# The magnitude of neonatal mortality and its predictors in Ethiopia: a systematic review and meta-analysis

**DOI:** 10.1101/626879

**Authors:** Yared Asmare, Wondimeneh Shibabaw, Tadesse Yirga, Abate Dargie, Tesfa Dejenie Hab-tewold

## Abstract

**Background:** Although neonatal death is a global burden, it is the highest in Sub Saharan Africa countries such as Ethiopia. This study was aimed to provide pooled national prevalence and predictors of neonatal mortality in Ethiopia.

**Objective:** To assess the pooled prevalence and predictors of neonatal mortality in Ethiopia.

**Search Strategy:** global databases were systematically explored. Systematically searched using the following databases: Boolean operator, Cochrane library, PubMed, EMBASE, HINARI, and Google Scholar. Selection, screening, reviewing and data extraction was done by two reviewers independently using Microsoft excel spread sheet. The modified Newcastle–Ottawa Scale (NOS) and the Joanna Briggs Institute Prevalence Critical Appraisal tools were used to assess the quality of evidence

**Selection criteria:** All studies conducted in Ethiopia and reporting the prevalence and predictors of neonatal mortality were included

**Data Collection and Analysis:** Data were extracted using a Microsoft Excel spreadsheet software and imported into STATA Version 14 s for further analysis. The pooled effect size with 95% confidence interval of neonatal mortality rate was determined using a weighted inverse variance random-effects model. Publication bias was checked using funnel plots, Egger’s and bagger’s regression test. Heterogeneity also checked by Higgins’s method. A random effects meta-analysis model was computed to estimate the pooled effect size (i.e. prevalence and odds ratio). Moreover, subgroup analysis based on region, sample size and study design were done.

**Results:** After reviewing 88 studies, 12 studies fulfilled the inclusion criteria and were included in the meta-analysis. The pooled national prevalence of neonatal mortality in Ethiopia was 16.3% (95% CI: 11.9, 20.7, I^2^ =88.6%). The subgroup analysis indicated that the highest prevalence was observed in Amhara region with a prevalence of 20.3% (95% CI: 9.6, 31.1, I^2^ =98.8) followed by Oromia, 18.8% (95%CI: 11.9,49.4, I^2^=99.5). Gestational age AOR,1.14 (95% CI: 0.94, 1.3), neonatal sepsis (OR:1.2(95% CI: 0.8, 1.5), respiratory distros (OR: 1.2(95% CI: 0.8, 1.5) and place of residency (OR:1.93 (95% CI:1.1,2.7) were the most important predictor.

**Conclusions:** neonatal mortality in Ethiopia was significantly decreased than the national report. There was evidence that neonatal sepsis, gestational age, respiratory distress were the significant predictors. We strongly recommended that health care workers should give a priority for the identified predictors.

## Background

The neonatal period is the first four weeks of a child’s life in which changes are very rapid and many critical events can occur in this period [1, 2]. Survival of new born babies had improved significantly through enhanced and specialized care. But, still it is the main reason of under-five death and risk of lifelong sequel [3–5]. Globally, the neonatal mortality rate (NMR) is declined from 49 % to 19%, but slower than under five mortality rate (dropped by 60%). Of all death of under-five, 40% were attributed with new born death [5] in which close to 1 and 2 million deaths occur on the day of birth, and in the first week of life respectively [6]. A review of 20 studies also indicated that the total NMR greatly varied between developed (4 to 46%) and developing (0.2 to 64.4%) countries [7]. Despite this, NMR shares the highest proportion of under-five mortality worldwide. [8–14]. According to the report of United Nations international child education and fund(UNICEF) and Ethiopian Demographic and Health Survey (EDHS), the neonatal mortality rate accounts 23% and 29%/1000 live births in Ethiopia [5, 15].

In Ethiopia, different studies have been conducted to assess the prevalence of neonatal mortality and associated factors. The findings of these uneven studies documented that there was a great variability in the prevalence of neonatal mortality across the regions of the country. Concerning predictors, these studies revealed that different maternal, neonatal and health service related factors influenced neonatal mortality; maternal residency [[16, 17]], history of antenatal, gestational age, neonatal sepsis, RDS, asphyxia were some of the factors associated with neonatal mortality [18, 19].Globally, there are different policies, strategies, and programs which work on prevention and care of neonate [20, 21]. However, it is sustained as the most cause of under-five mortality world-wide [11, 22-25].Despite different strategies, health policies (like HSDP IV 2010-2015) and interventions were implemented to prevent and care neonatal death, the rate of neonatal mortality is high. Additionally, Ethiopia achieved millennium development goal in reducing child mortality by two-third, but neonatal mortality is still high. Several studies were conducted and showed important results, including prevalence of mortality and its major predictors in the study area. However, there is variation in the prevalence and associated factors of studies based on region, sample size and the study design that have been used.as far as our knowledge, there has been no any systematic review and meta-analysis of studies reporting on the neonatal mortality and its predictors. So that we believe synthesizing these studies may fill the gap in the literature and provide stronger evidence for policy-making as there is an increasing recognition of systematic review and meta-analysis findings in the policy-making process. Thus, further meta-analysis and synthesis of the available evidence is needed now. Hence, this study aimed to synthesize nationally available evidence on the prevalence of neonatal mortality and the association with different variables in Ethiopia. The findings of this study will be used as an input to policy makers and program planners working in the area of neonatal health.

## Methods

### Searching strategies

The PRISMA guidelines protocol was used to write the systematic review [22]. The reviewer follows PRISMA systematic review protocol as a reporting guideline (for the PRISMA checklist, eligible studies for the study were selected in terms of titles alone, abstracts, and then full-text articles, based on inclusion criteria. Cochrane library, PubMed, EMBASE, HINARI, and Google Scholar was systematically searched for articles. The studies were accessed using the following search terms: “neonatal mortality”,” predictors”, “neonatal death”, “newborn”, “prevalence”, “neonatal mortality” and” Ethiopia”. The search terms were used individually and in combination using “AND” and “OR” Boolean operators. In addition, after identification of included studies, cross-references were searched to identify more eligible studies. The search was guided by PECO: Population - neonates (age < 28 days); occurrence of death within 28 days after delivery.

### Inclusion and exclusion criteria

Studies with the following major criteria were considered for inclusion. Observational (i.e. cross-sectional, case–control and cohort) studies in Ethiopia, which reports the prevalence and predictors of neonatal mortality were included. Articles published in English language was considered as further inclusion criteria. On the other hand, studies which did not report the outcome and articles without full-text were excluded. Corresponding authors were approached by email at least twice to access full-texts.

### Screening and data extraction

Two reviewers (YA and WS) screened titles and abstracts against the inclusion criteria-. Then, the full-texts of articles were accessed and independent assessment was carried out by two reviewers based on the predetermined inclusion and exclusion criteria. Discrepancies between the reviewers were resolved through discussion and common consensus of all investigators. Data were extracted from the included papers by AD, TD and TY independently extracted data from a random sample of 20% of papers to check consistency; consequently, there was no differences.

### Assessment of study quality

Structured data abstraction form was constructed in Microsoft excel. In each abstract and full text of the article, which was considered to be appropriate; a special emphasis was given for clearness of objective, data about the study area, study design, year of publication, study population, sample, size respondent rate, prevalence/incidence of neonatal death and other useful variables were recorded (table 1). The Joanna Briggs Institute Prevalence Critical Appraisal Tool for use in systematic review for prevalence study was used for critical appraisal of studies [26]. Moreover, methodological and other quality of each article was assessed based on a modified version of the Newcastle-Ottawa Scale for cross-sectional study adapted from Modesti et al[27].

**Table 1:**
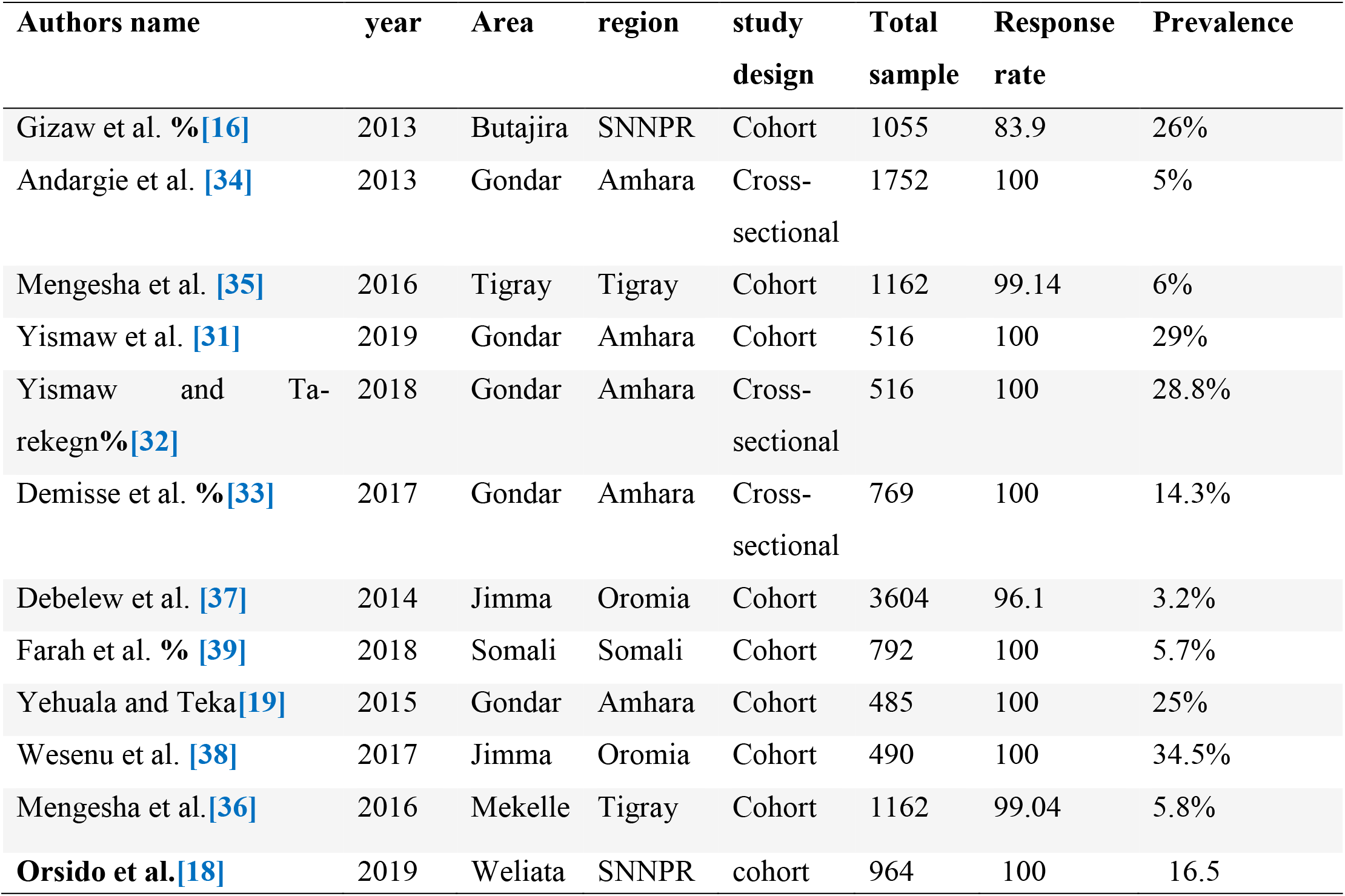
Descriptive summary of 12 studies included in the meta-analysis of the prevalence of neonatal mortality.

| Authors name | year | Area | region | study design | Total sample | Response rate | Prevalence |
| --- | --- | --- | --- | --- | --- | --- | --- |
| Gizaw et al. %[16] | 2013 | Butajira | SNNPR | Cohort | 1055 | 83.9 | 26% |
| Andargie et al. [34] | 2013 | Gondar | Amhara | Cross-sectional | 1752 | 100 | 5% |
| Mengesha et al. [35] | 2016 | Tigray | Tigray | Cohort | 1162 | 99.14 | 6% |
| Yismaw et al. [31] | 2019 | Gondar | Amhara | Cohort | 516 | 100 | 29% |
| Yismaw and Ta-rekegn%[32] | 2018 | Gondar | Amhara | Cross-sectional | 516 | 100 | 28.8% |
| Demisse et al. %[33] | 2017 | Gondar | Amhara | Cross-sectional | 769 | 100 | 14.3% |
| Debelew et al. [37] | 2014 | Jimma | Oromia | Cohort | 3604 | 96.1 | 3.2% |
| Farah et al. % [39] | 2018 | Somali | Somali | Cohort | 792 | 100 | 5.7% |
| Yehuala and Teka[19] | 2015 | Gondar | Amhara | Cohort | 485 | 100 | 25% |
| Wesenu et al. [38] | 2017 | Jimma | Oromia | Cohort | 490 | 100 | 34.5% |
| Mengesha et al.[36] | 2016 | Mekelle | Tigray | Cohort | 1162 | 99.04 | 5.8% |
| **Orsido et al.[18]** | 2019 | Weliata | SNNPR | cohort | 964 | 100 | 16.5 |

### Data synthesis and statistical analysis

Data were extracted using a Microsoft Excel spreadsheet software and imported into STATA Version 14 software for further analysis. The pooled effect size with 95% confidence interval of national neonatal mortality rate was determined using a weighted inverse variance random-effects model [28]. Heterogeneity across the studies was assessed using I^2^ statistic where 25, 50 and 75% representing low, moderate and high heterogeneity respectively[29]. A Funnel plot, beggar’s and Egger’s regression test were used to check publication bias[30]. Moreover, subgroup analysis based on study area (region), study design and sample size were done. Log odds ratios were used to examine the association between mortality and its major predictors.

## Results

### Characteristics of included studies

The abstract search resulted in 136985 PUBMED references when we search with” neonatal morality) OR Ethiopia in which the report is not limited to the study area. When farther advanced searches are attempted the search, strategy retrieved 88 potential articles, of which 12 full text articles that fulfill the eligible criteria with a total sample size of 12397 neonates were included in the final analysis for the current systematic review and meta-analysis (Fig. 1). Information about authors, publication year, population, study area, region, study design, outcome and main results from the selected articles were extracted and summarize on table. the overall respondent rate was between eighty-three to hundred percent.

**Fig. 1:**
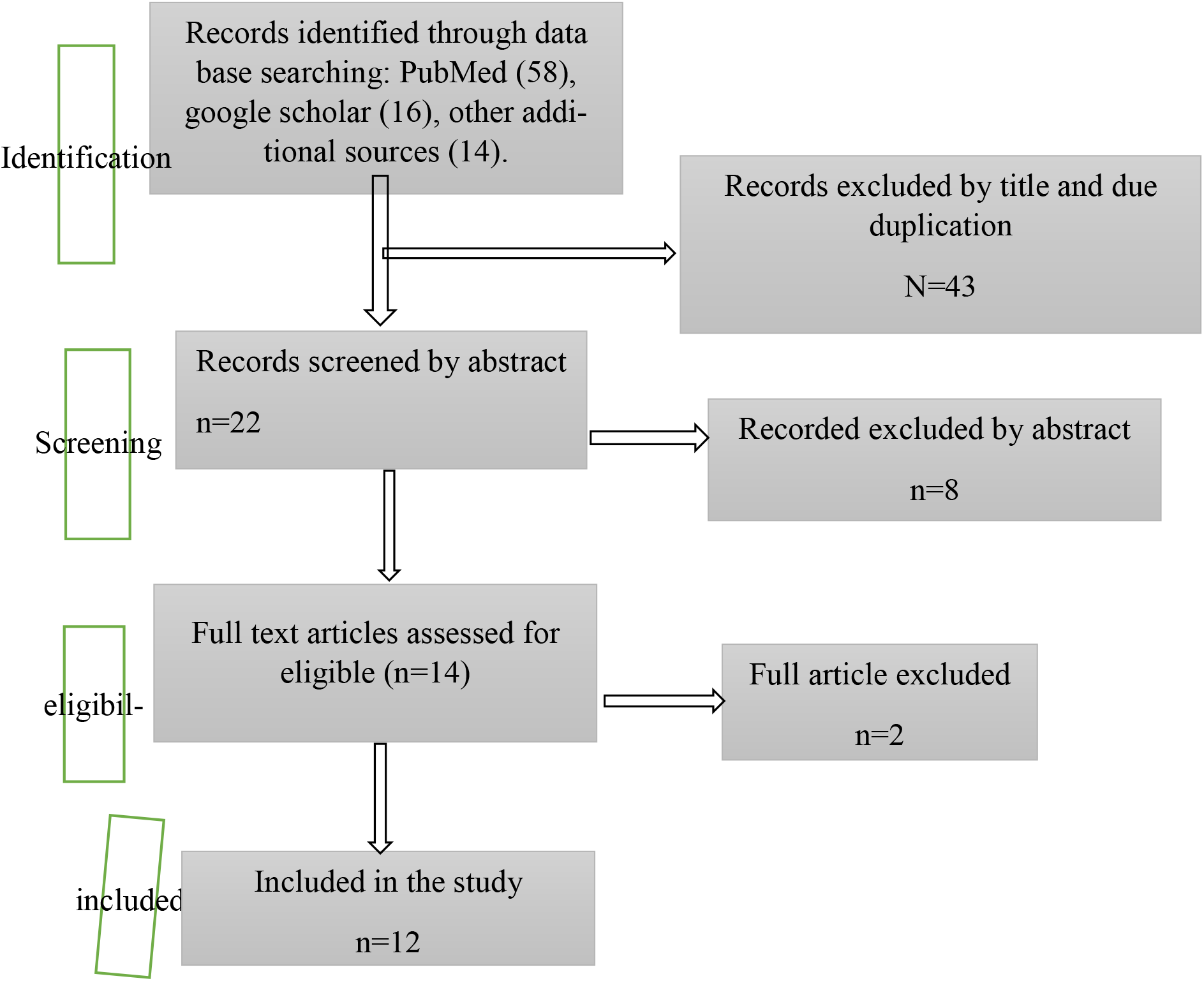
PRISMA flow diagram for showing screening and selection process of studies.

#### Features of Studies

All studies were done in Ethiopia and published in an indexed journal. Eight of them were cohort (both retrospective and prospective) and the remaining three were cross sectional studies. Detailed screening and selection process were shown in Figure 1.

### Study characteristics

The studies were conducted in Amhara [19, 31-34], Tigray [35, 36], Southern Nations, Nationalities and Peoples (SNNP) [16], Oromia[37, 38] and Somali [39] region. Eight were cohort and three were cross sectional studies. The sample size of the studies ranging from 485 to 3,604.

### prevalence of neonatal mortality

In the current systematic review and meta-analysis, the pooled prevalence estimates of neonatal mortality were described by forest plot (Fig. 2). The pooled prevalence of neonatal mortality from the random effect’s method was found to be 16.3% (95% CI; 12.1–20.6).

**Figure 2:**
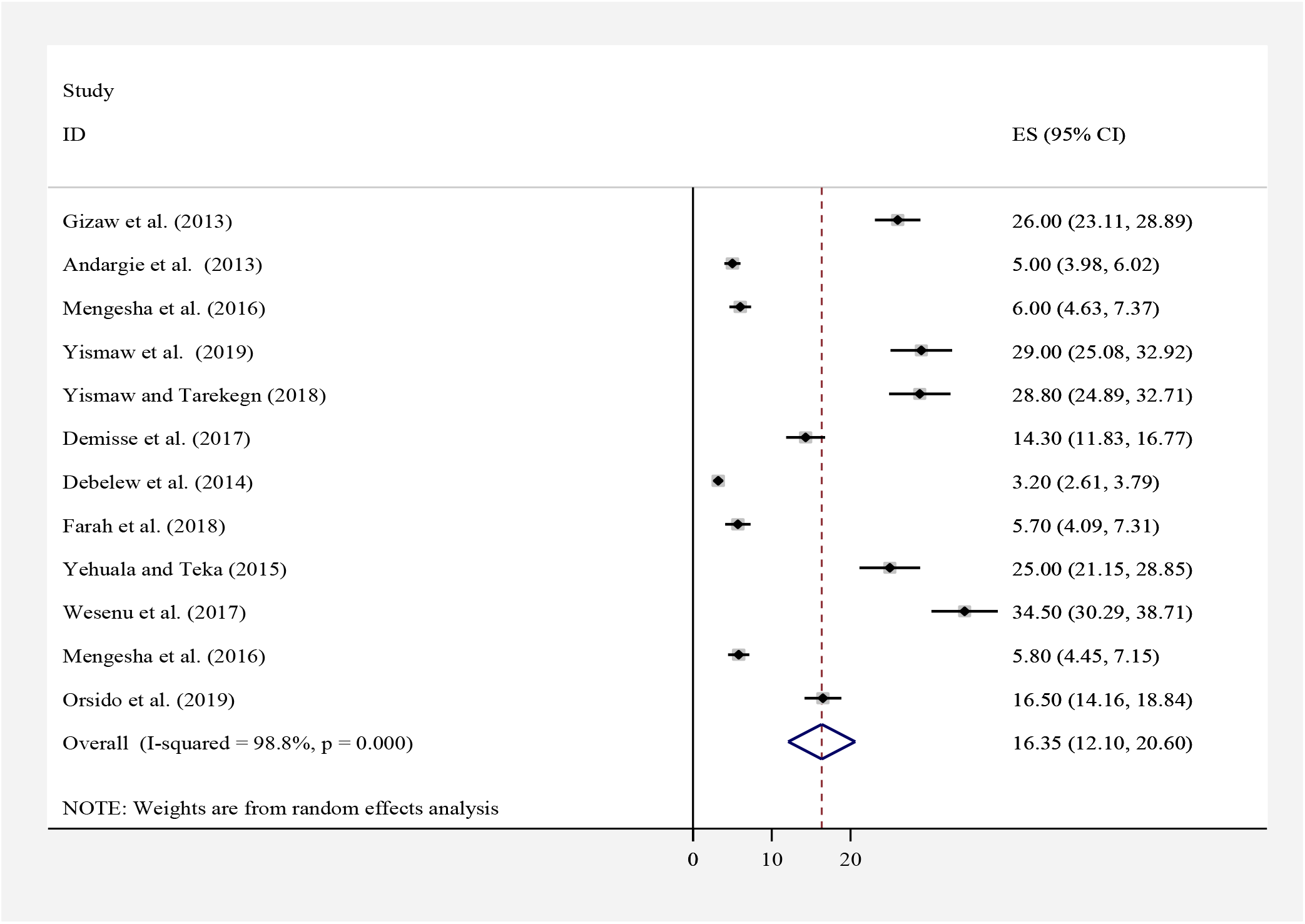
Forest plot of the pooled prevalence of neonatal mortality.

### Investigation of heterogeneity and publication bias

Consequently, the analysis showed a substantial heterogeneity of egger test (p < 0.001) and I^2^ statistics (I^2^ = 83%) for neonatal mortality. The discrepancy in the pooled estimates of the prevalence was adjusted through subgroup analysis based on the region, where the study conducted and based on the types of study design. A sensitivity analysis was conducted to check the stability of summary estimate. The funnel plot was found to be asymmetric (fig 3) and Egger’s and bagger’s test was found to be significant publication bias at p value ≤ 0.001 which confirms that there is a publication bias.

**Fig.3:**
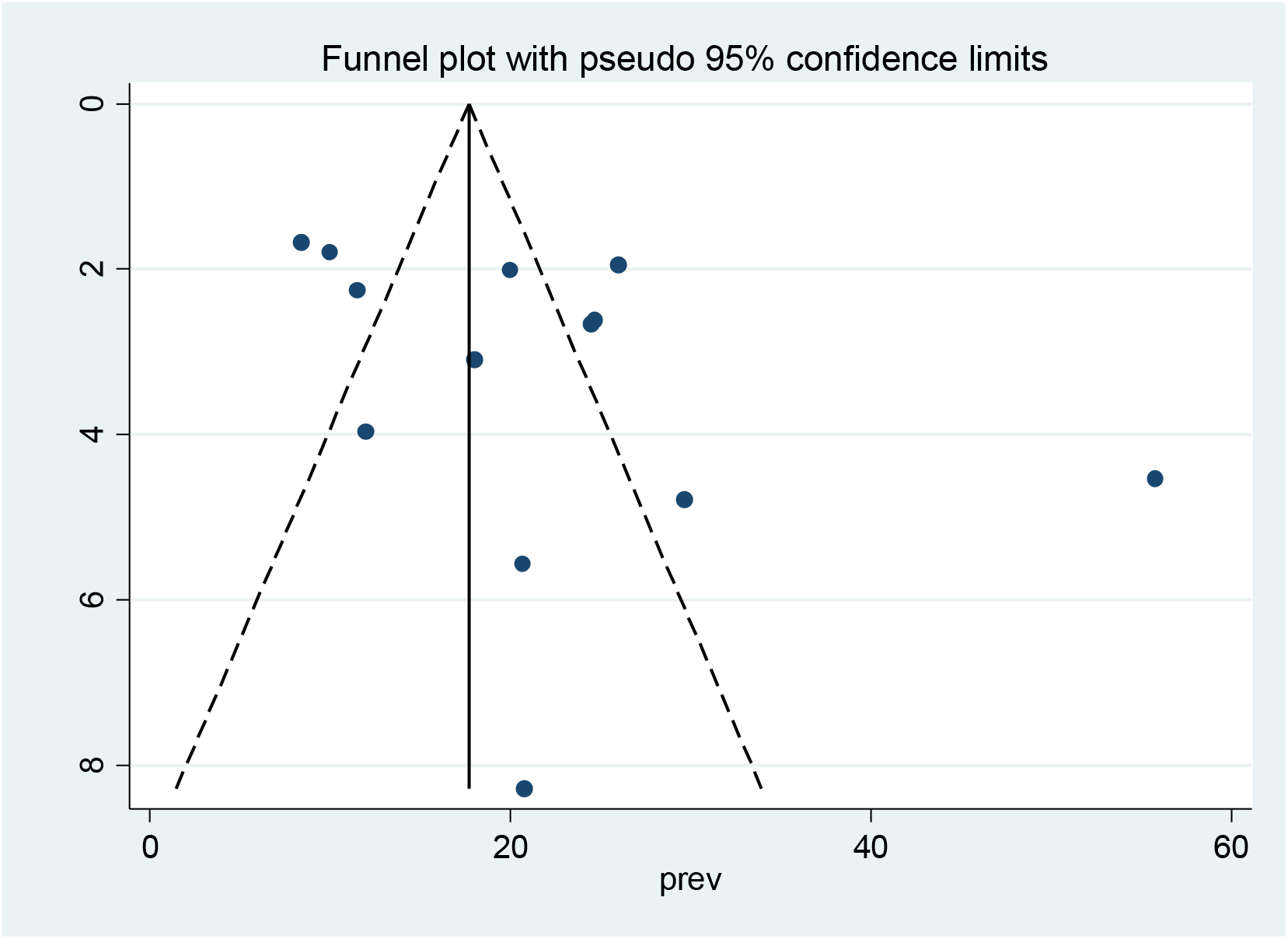
Funnel plot to show the distribution of 12 studies.

### Meta-regression analysis

Based on the subgroup-analysis result, the highest prevalence (20.3%; 95% CI:9.6 −31.1) was reported in Amhara region with regard to sample size, the prevalence of neonatal mortality was higher in studies having a sample size < 800, 22.7% (95%CI:12.8, 32.7) compared to those having a sample size ≥ 800,8.9 (95%CI:4.9, 12.9) (Table 3).

**Table 3:** Meta-regression analysis of neonatal sepsis with heterogeneity of neonatal mortality.

| Heterogeneity source | Coefficients | Std. Err. | P-value |
| --- | --- | --- | --- |
| Publication Year | **-1.243** | **4.738** | **0.64** |
| Sample size | **0.01800** | **0.290** | **0.86** |

## Predictors of neonatal mortality

### Gestational age

Seven studies[17, 31, 33, 36-38, 40], examined the association between gestational age and neonatal mortality. The pooled odds ratio was 1.32 (95% CI: 1.07, 1.58), *I*^2^= 49%). Neonates born as term had 14% lower chance of death compared with preterm neonates (Fig. 4). Begg’s (p= 0.37) and Egger’s (p=0.7) tests did not reveal significant publication bias.

**Figure 4:**
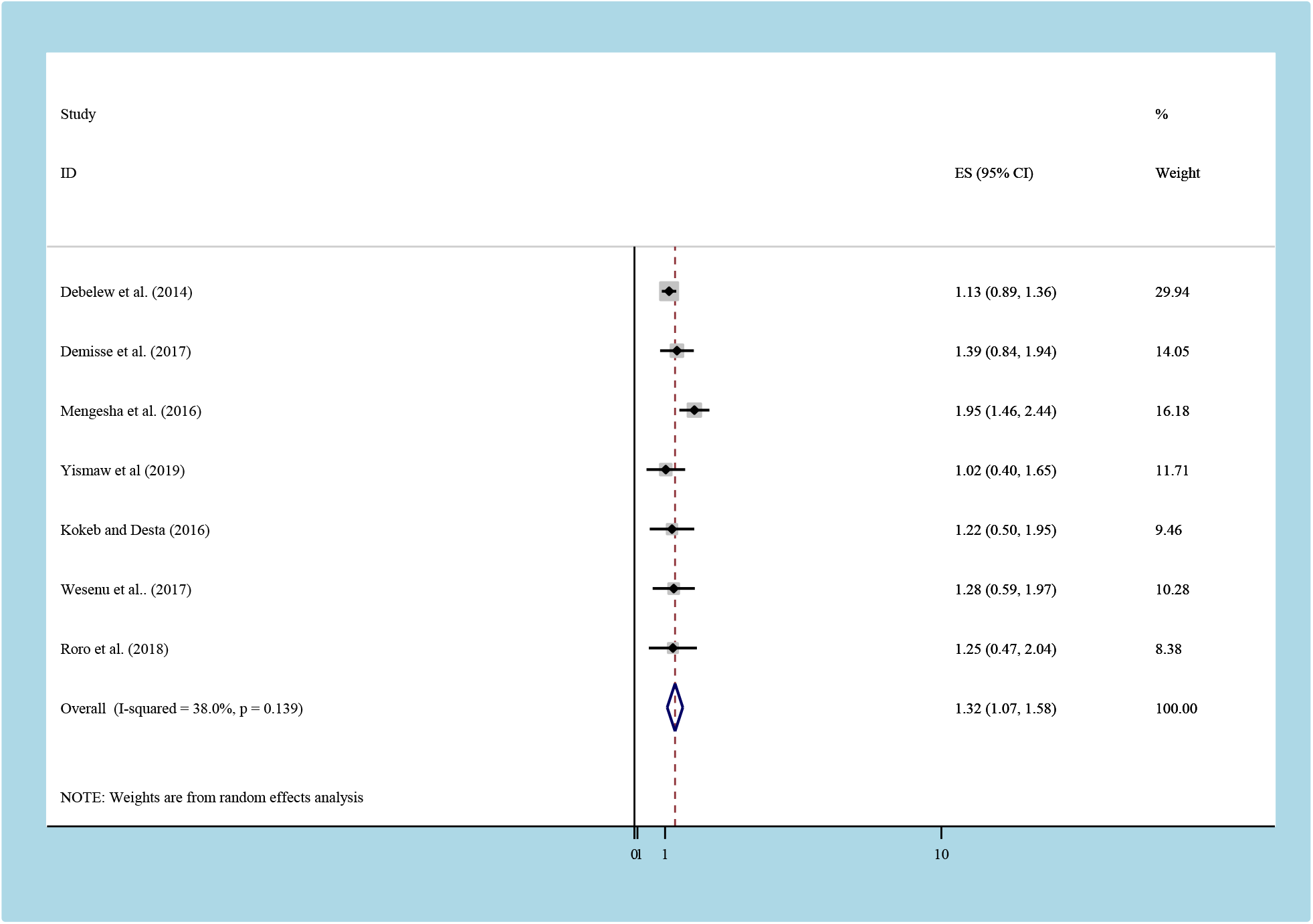
the pooled odds ratio of the association between GA and neonatal mortality.

### Residency

Six studies[31-33, 36, 37, 41] reported the association between place of residency and neonatal mortality. The pooled odds ratio was1.93 (95% CI:1.1,2.7; *I*2= 3%) (fig 5). Neonates who were born in rural area were found to be 98% higher risk of mortality than their counterpart although not statistically significant. Begg’s (p=0.99) and Egger’s tests (p=0.663) showed that there was no significant publication bias.

**Figure 5:**
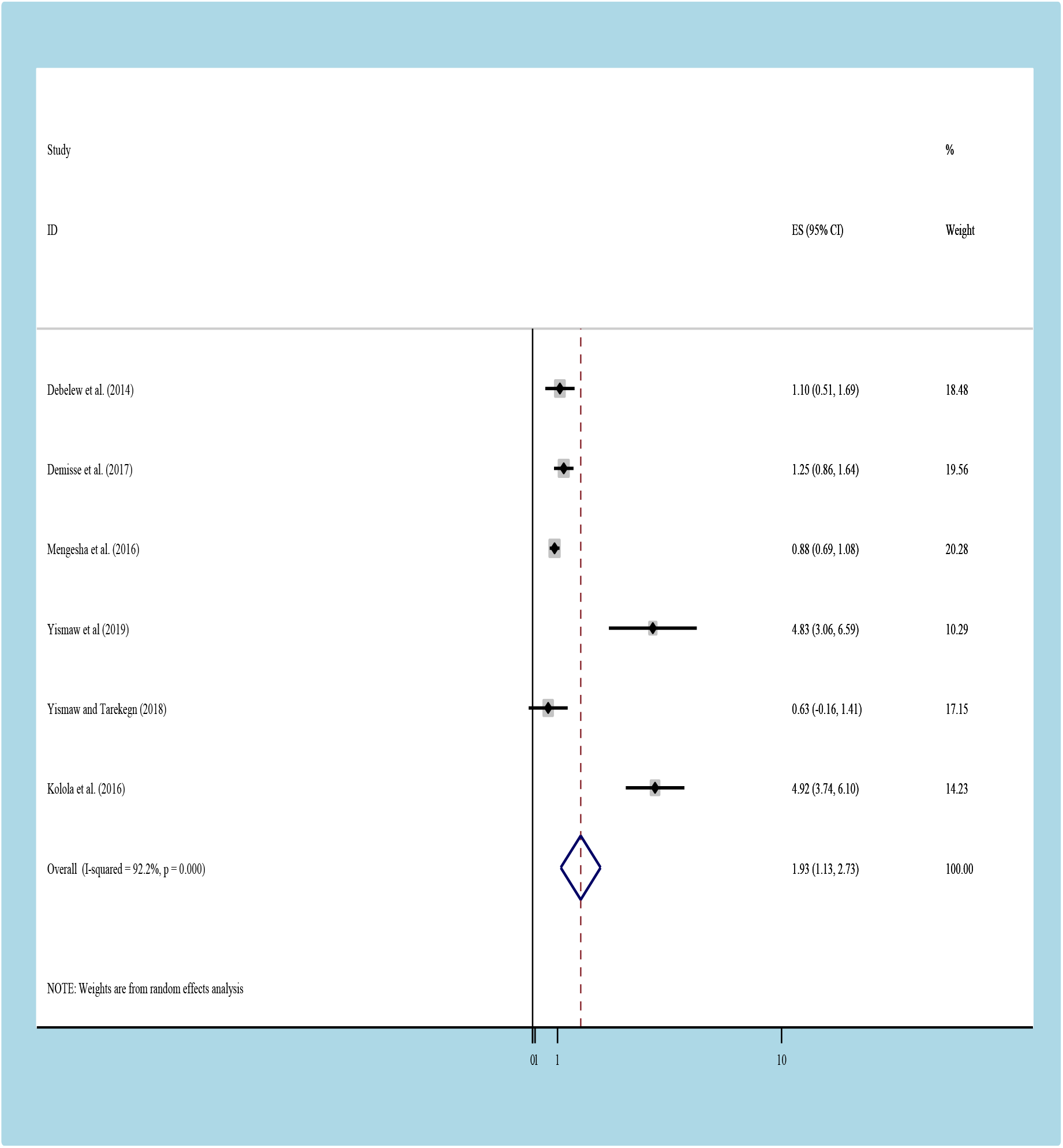
the pooled odds ratio of the association between residency and neonatal mortality.

### Respiratory distros syndromes

According to these finding, those neonates having RDS were also associated with neonatal mortality (fig 6). Neonates who had RDS were nearly one and half times more likely to die as compared to those who did not have RD (OR= 1.2,95% CI: 0.8, 1.5). We observed that a no any heterogeneity across the studies (I^2=^ 4.43% (d.f. = 5) with no any publication bias with begger (p = 1.000) and egger (p= 0.138) tests.

**Figure 6:**
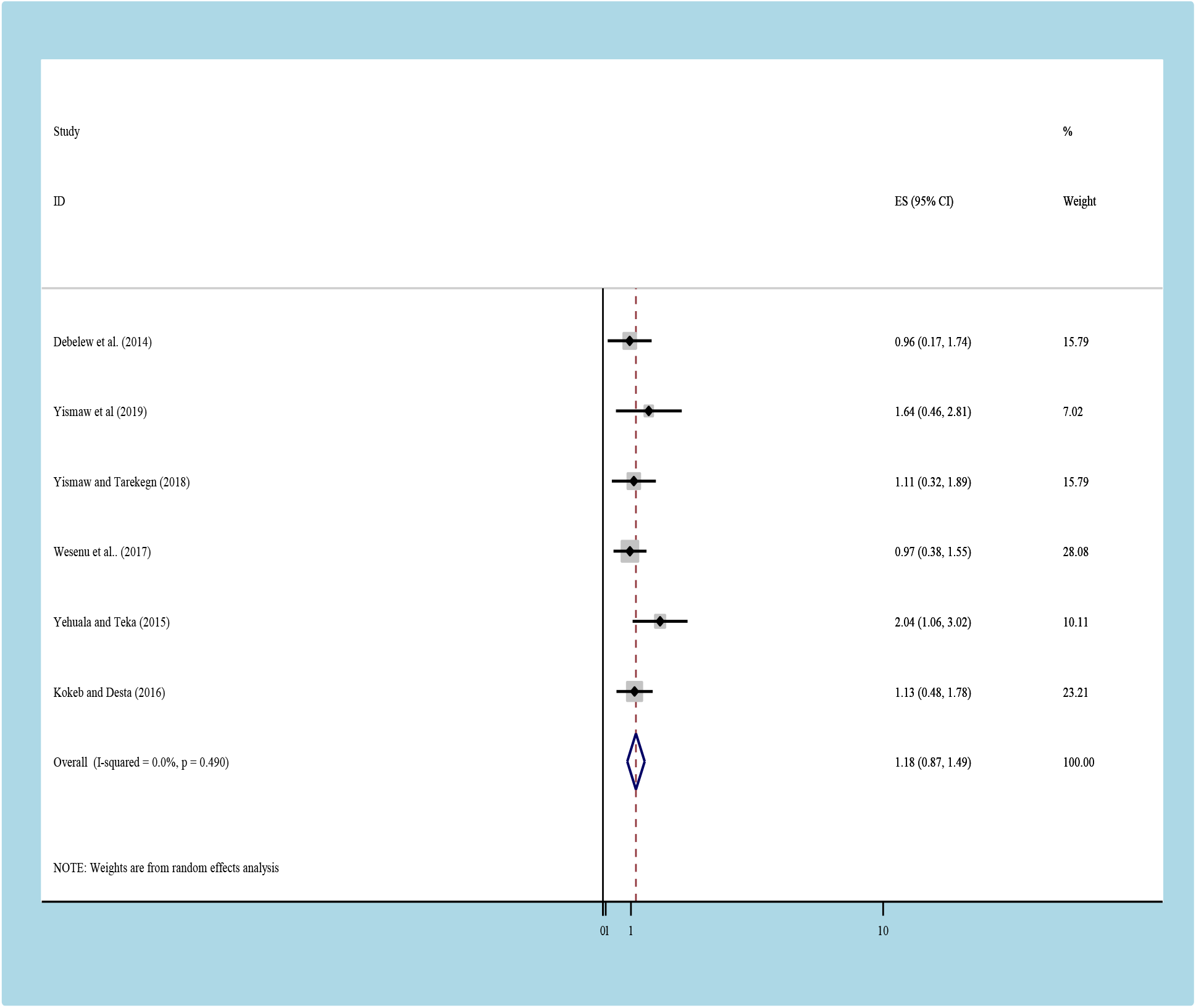
the pooled odds ratio of the association between RDS and neonatal mortality.

### Neonatal sepsis

To assess the association of neonatal sepsis with neonatal mortality, we have include five studies[33, 36-38, 41], Patients who had neonatal sepsis had almost one and half times higher chance of neonatal death compared to those patients without sepsis OR: 1.23(95% CI:1.1, 1.4) (Figure 7). The heterogeneity test showed extreme evidence of variation across studies (I^2^= 90%, P= 0.001. Moreover, the result of Egger’s test to examine publication bias showed no statistically significant evidence of publication bias (begge, P= 0.806) and (egger P=0.511).

**Figure 7:**
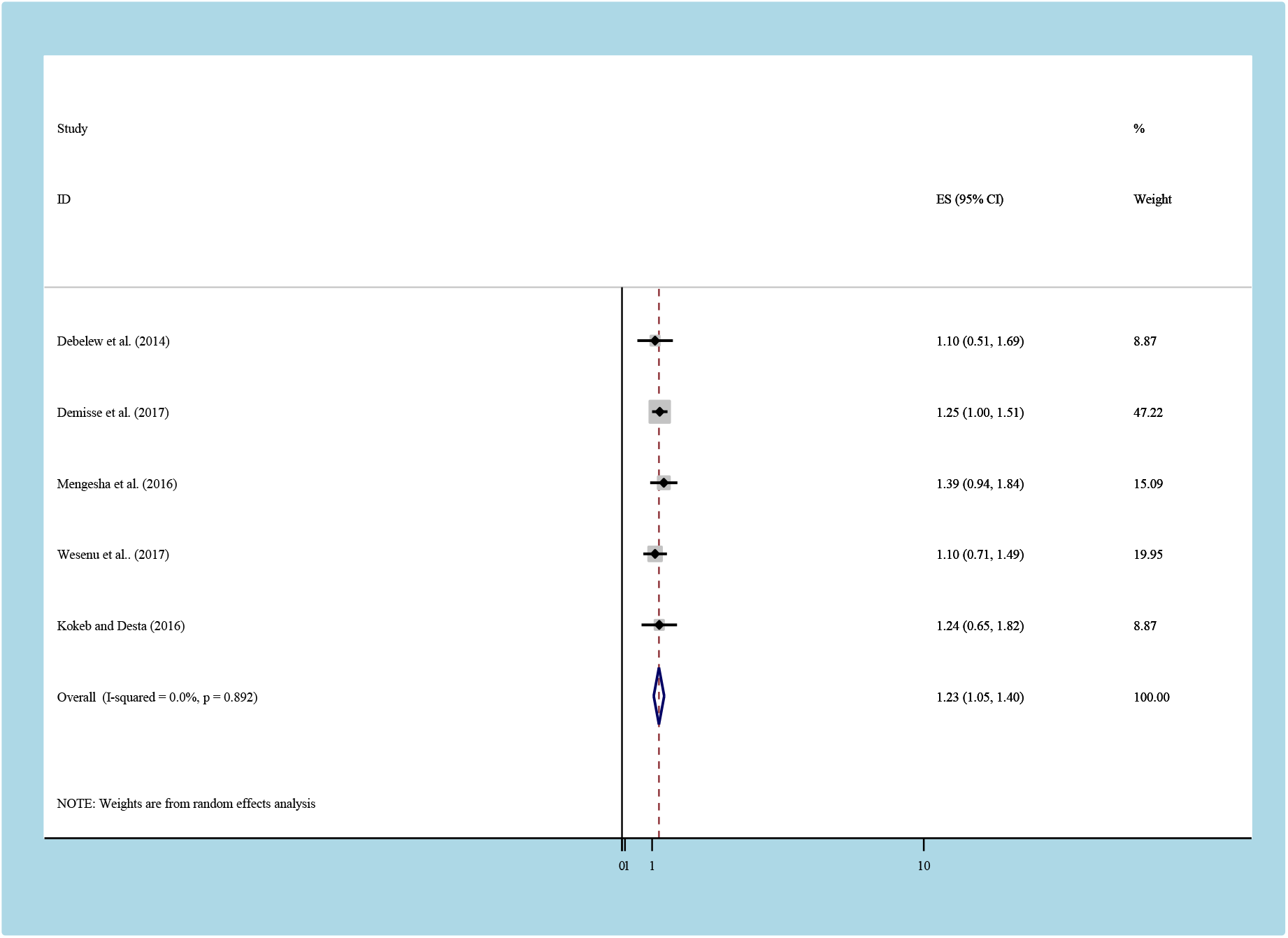
Pooled odds ratio of the association between neonatal sepsis and neonatal mortality.

## Discussion

This systematic review and meta-analysis revealed the national burden of neonatal mortality. We also sought, gestational age, neonatal sepsis and residence significantly predicted neonatal mortality but not respiratory distress syndrome. Our study showed that the national prevalence of neonatal mortality was 16.3%. This is in line with a study findings in Cameroon, South Africa, South Sudan and Mauritanian [15, 42] [40]. However, our finding was lower than the UNICEF national 2016 report[3, 5] and other national reports in Africa[43, 44], Europe, USA, Central and West Asia [45–47]. This marked difference might be attributed to difference in methodology, sample size, study period and geographic area. For example, the UNICEF report includes all regions and cities of Ethiopia but the current study includes studies only from five regions. Additionally, some of the previous reports limited study area instead of nationwide reports. Moreover, the difference in the study period could be a relevant factor as standard or care and treatment modalities changes over time. Additionally, though, neonatal health care services in Ethiopia have been remaining less consolidated, systematic efforts have been gaining momentum since relatively recent. Furthermore, the NICU service in our countries is not well included Level three (subspecialty) NICU all over the countries.

The subgroup analysis indicated that the highest prevalence of mortality was observed in Amhara region (20.3%) whereas the lowest prevalence was observed in Somalia (5.7%). The possible reason might be that in Amhara region five studies were included compared with other regions. Additionally, the sampled population included in Amhara region were higher than other regions. These studies reported that, having neonatal sepsis was significantly increased the risk of mortality. Those who had neonatal sepsis were nearly two times increase risk of mortality as compared to those who did not have sepsis. This finding is supported by a result in developed and developing country[19] [48] [7], [49], [50] [51], ([52], [53]. The possible reason might be due to newborns have many physiologic challenges when adapting to the extrauterine environment which might contributed to common problem like immature immunity, RDS., neurologic, cardiovascular, hematologic, nutritional, gastrointestinal and poor thermoregulation which farther increase risk of sepsis and mortality. We also found that neonates living in rural areas were more vulnerable to death than their urban counterparts. Neonates from rural households were nearly two times more likely to die as compared to their counterparts. This finding is similar to a study conducted in Washington, Louisiana, and Tennessee [54].This variance could be due to due to the fact that rural residents are still relatively disadvantaged in terms of infrastructures, knowledge and awareness difference, distance from the site. Another possible reason could be that people living in rural areas tend to be poorer than their urban counterparts’ area, a factor known to have an impact on the neonatal outcome.GA is also another important determinant of neonatal mortality in our meta-analysis. Accordingly, neonates born as preterm were almost one and half times more likely to die than those term neonates [49-57][55],[56], [52], [53],[50], [7] [51], [49, 57].RDS are an important determinant of neonatal mortality although not statistically significant in our meta-analysis. Neonates who had RDS were one times more likely to die as compared to those who did not have RD. Consistent results have been recorded in our countries and other studies [32,34,38,45]. This might be due to that Babies with RDS don’t have a protein called surfactant that keeps small air sacs in the lungs from collapsing which increase the risk of neonatal mortality

This systematic review and meta-analysis were conducted to assess the pooled prevalence of neonatal mortality and its predictors in Ethiopia. Conducting this type of study will act as an input for program planners and policy makers working in the area of neonatal care and also indicates the quality of health care and the welfare of the society. This study is the first meta-analysis in the study area with novel findings. Even though this analysis has delivered valued evidence regarding the level of neonatal mortality and its predictors, there were some limitations, which we address below: This meta-analysis represented only studies reported from five regions of the country. Therefore, the regions may be under-represented due to the limited number of studies included.

## Conclusion

Despite the variation across regions, in Ethiopia, at least one out of ten newborn had risk of death before their first birth date. Neonatal sepsis and residence were an important predictor of neonatal mortality. Therefore, based our findings, we recommend particular emphasis shall be given to the rural communities. Additionally, the government should have to strengthen any service related with reducing the neonatal mortality like expanding NICU all over the country and particular emphasis should be given for those neonate’s diagnosis as preterm birth, neonatal sepsis and RDS.

## Abbreviations

CI: Confidence Interval
OR: Odds Ratio
PRISMA: Preferred Reporting Items for Systematic Reviews and Meta-Analyses
RDS: respiratory distros syndromes: Southern Nations, Nationalities, and Peoples
WHO: World Health Organization

## Declarations

### Authors’ contributions

YA Conception of research protocol, study design, literature review, data extraction, data analysis, interpretation and drafting the manuscript. WS, AD, TD and TY: data extraction, quality assessment, data analysis and reviewing the manuscript. All authors have read and approved the manuscript.

### Ethics approval and consent to participate

Not applicable.

### Availability of data and materials

The data analyzed during the current systematic review and meta-analysis is available from the corresponding author on reasonable request.

### Consent for publication

Not applicable.

